# LIS1 determines cleavage plane positioning by regulating actomyosin-mediated cell membrane contractility

**DOI:** 10.1101/751958

**Authors:** Hyang Mi Moon, Simon Hippenmeyer, Liqun Luo, Anthony Wynshaw-Boris

## Abstract

Heterozygous loss of human *PAFAH1B1* (coding for LIS1) results in the disruption of neurogenesis and neuronal migration via dysregulation of microtubule (MT) stability and dynein motor function/localization that alters mitotic spindle orientation, chromosomal segregation, and nuclear migration. Recently, human induced pluripotent stem cell (iPSC) models revealed an important role for LIS1 in controlling the length of terminal cell divisions of outer radial glial (oRG) progenitors, suggesting cellular functions of LIS1 in regulating neural progenitor cell (NPC) daughter cell separation. Here we examined the late mitotic stages NPCs *in vivo* and mouse embryonic fibroblasts (MEFs) *in vitro* from *Lis1*-deficient mutants. *Lis1*-deficient neocortical NPCs and MEFs similarly exhibited cleavage plane displacement with mislocalization of furrow-associated markers, associated with actomyosin dysfunction and cell membrane hyper-contractility. Thus, it suggests LIS1 acts as a key molecular link connecting MTs/dynein and actomyosin, ensuring that cell membrane contractility is tightly controlled to execute proper daughter cell separation.

## INTRODUCTION

Human *Lissencephaly-1* (*LIS1*) haploinsufficiency is the most common genetic cause of a neuronal migration disorder called lissencephaly (Dobyns et al., 1993; Hattori et al., 1994; Moon et al., 2013). LIS1 is a key regulator of cytoplasmic dynein and microtubules (MTs), and it has been proposed as an important molecular link coordinating both MTs and actin (Kholmanskiki 2003; Kholmanskiki 2006; Jheng et al., 2018). Besides an indispensible role for LIS1 in neuronal migration in post-mitotic neurons, *Lis1* mutant mouse studies suggest pivotal roles of LIS1 in neocortical neural progenitor cell (NPC) division (Yingling et al., 2008; Youn et al., 2009; Hippenmeyer et al., 2010; Bershteyn et al., 2017) by regulating mitotic spindles (Yingling et al., 2008; Moon et al., 2014), consistent with other mutants of MT/dynein-associated proteins such as LGN, NDE1, and NDEL1 (Fuga et al., 2004; Bradshaw and Hayashi 2017; Doobin et al., 2016; Wynne et al., 2018). However, unlike these other mutants, *Lis1* mutants specifically displayed a decrease in neuroepithelial stem cells in the neocortex and subsequent neonatal death compared with a less catastrophic phenotype seen in radial glial (RG) progenitors (Yingling et al., 2008). Our recent studies with human induced pluripotent stem cells (iPSCs) of Miller-Dieker syndrome, a severe form of lissencephaly caused by heterozygosity of more than 20 genes including *LIS1*, demonstrated a prolongation of mitotic division time of outer radial glial (oRG) progenitors but not ventricular zone radial glial (vRG) progenitors (Bershteyn et al., 2017), suggesting that additional cellular functions of LIS1, other than MT/dynein-mediated mitotic spindle regulation, dictate further mitotic progression from mitotic (M) phase to anaphase/telophase in late stages of mitosis and daughter cell separation.

During late stages of mitosis, microtubules (MTs) play a central role to provide important positional cues for the cleavage furrow localization at the equatorial cortex by inducing the assembly of the central spindle (Bement et al., 2005; von Dassow, 2009; Werner et al., 2007). In addition, crosstalk between astral MTs and cortical actin generates inhibitory signals at the polar cortex that restrict cleavage furrow positioning to the equatorial cortex (Canman et al., 2003; Foe and von Dassow, 2008; Murthy and Wadsworth, 2008). At the molecular level, cleavage plane positioning is regulated by a small GTPase RhoA (Bement et al., 2005; Nishimura and Yonemura, 2006) that mediates recruitment of contractile ring components to the equatorial cortex, including the scaffold protein Anillin (D’Avino et al., 2008; Gregory et al., 2008; Oegema et al., 2000; Piekny and Glotzer, 2008) and filamentous Septin (Kinoshita et al., 2002; Maddox et al., 2007; Oegema et al., 2000). Similarly, Myosin II is accumulated at the equatorial cortex and serves as a force generator cooperated with filamentous actin (F-actin) (Glotzer, 2005; Straight et al., 2003; Zhou and Wang, 2008). When fewer or elongated astral MTs destabilize the cleavage furrow, MT dysfunction induces the dysregulation of actomyosin–mediated cell membrane contractility, RhoA and F-actin mislocalization outside of the midzone/cleavage furrow, cell shape oscillation, and subsequent failure in completion of mitosis (Canman et al., 2003; Murthy and Wadsworth, 2008; Rankin and Wordeman, 2010; Watanabe et al., 2008).

However, it is still unknown whether LIS1 particularly is involved in spatiotemporal regulation of actomyosin activity and cell membrane contractility during late stages of mitosis and cytokinetic completion step when daughter cell separation is crucial. We hypothesized that the LIS1-dynein-MT network may be essential for timely progression of late stages of mitosis such as cytokinesis, perhaps by fine-tuning of actomyosin and cell membrane contractility. Therefore, we assessed the phenotypes of neocortical NPCs in embryonic neocortical development from *Lis1*-deficient mice, and monitored the entire duration of mitotic events of *Lis1*-deficient mouse embryonic fibroblasts (MEFs), using time-lapse live cell imaging. we found defects in cleavage plane positioning and dysregulation of cell membrane contractility in *Lis1*-deficient MEFs. Here we demonstrate that LIS1 determines cleavage plane positioning through a RhoA-actomyosin signaling pathway, defining a novel molecular mechanism of LIS1 underlying spatiotemporal regulation of cell membrane contractility during mitosis.

## RESULTS

### Mislocalization of RhoA and Anillin in *Lis1* mutant neocortical neural progenitor cells (NPCs)

To elucidate molecular mechanisms underlying LIS1-dependent NPC regulation during neocortical development, mitotic phenotypes of *Lis1*-deficient NPCs were analyzed by conducting genetic and immunohistochemical approaches on mouse neocortices derived at embryonic day 14.5 (E14.5) (**Fig. 1A-E**). We took advantage of mosaic analysis with double markers on chromosome 11 mouse line (MADM-11; Hippenmeyer et al., 2010), since the *Lis1* gene is located on chromosome 11 away from the centromere. To deplete *Lis1* sparsely in neocortical NPCs during early embryonic development, we first generated *MADM-11^TG/TG,Lis1^* (TG: tdTomato-GFP fusion) mice co-expressing the heterozygous *Lis1* knock-out (KO) allele. These mice were mated with *MADM-11^GT/GT^;Emx1-Cre^Cre/+^* (GT: GFP-tdTomato fusion) to generate the experimental mosaic animals which carry sparsely labeled NPCs with different expression levels of LIS1 (*MADM-11^GT/TG,Lis1^;Emx1-Cre^Cre/+^*, abbreviated as *Lis1-MADM*) (**Fig. 1 B**). In this line, we identified three distinct subpopulations of labeled NPCs with different LIS doses; *Lis1^+/+^* (red, labeled with tdTomato, 100% LIS1 wild-type (WT) levels), *Lis1^ko/+^* (yellow, double positive for GFP and tdTomato, 50% LIS1 WT levels), and *Lis1^ko/ko^* (green, labeled with GFP, 0% LIS1) NPCs in an unlabeled *Lis1^ko/+^* heterozygous background. The fluorescence of each cell enabled us to distinguish the genotype of each cell. The same mating scheme was used to generate WT control animals (*MADM-11^GT/TG^;Emx1-Cre^Cre/+^*, abbreviated as *WT-MADM*) where green, red, and yellow cells are all WT expressing normal levels of 100% LIS1.

**Fig. 1.**
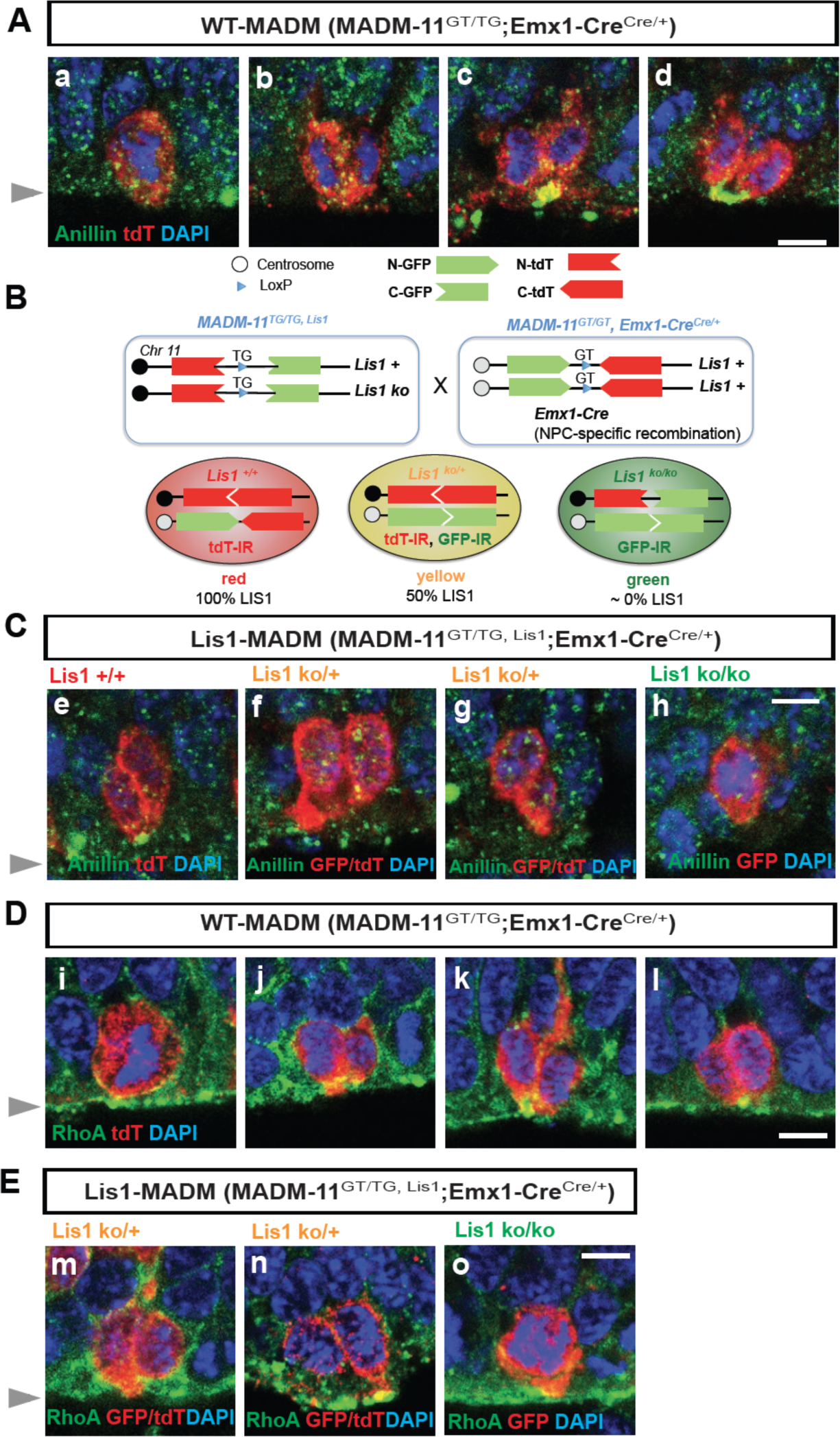
Cleavage plane formation and positioning in the neocortical neural progenitor cells (NPCs) in *WT-MADM* and *Lis1-MADM* embryos. **(A)** wild-type (WT) NPCs displayed recruitment of Anillin to the basal equatorial cortex and ultimately the Anillin-ring moved to the apical surface of the ventricular zone, forming a ‘U’-like shape. **(B)** Schematic representation of *Lis1-MADM* mating scheme and three types of neocortical NPCs with different LIS1 expression levels. Immunoreactivity (IR) from immunohistochemistry experiment with anti-GFP and anti-tdT-c-Myc antibodies was indicated. **(C) e.** Midbody-associated Anillin localization in WT NPCs (*Lis1^+/+^*), **f,g.** Midbody-associated Anillin distribution was not detected in *Lis1* heterozygous NPCs (*Lis1^ko/+^*). **h.** Complete knock-out (KO) of LIS1 (*Lis1^ko/ko^*) in NPCs results in mitotic arrest at the prometaphase- or metaphase-like time-points. **(D) i,j.** WT NPCs displayed apical membrane-associated RhoA during cytokinesis. **k.l.** RhoA was also co-localized with the midbody, at the future cleavage furrow. **(E) m.n.** The cytosol from only one daughter cell retained RhoA-positive puncta in *Lis1* heterozygous NPCs (*Lis1^ko/+^*). **o.** Complete KO of LIS1 (*Lis1^ko/ko^*) resulted in detachment of NPCs away from the RhoA-positive apical membrane. Grey arrowheads: ventricular surface (apical). Scale bars: 5 μm.

To determine whether *Lis1* deficiency in neocortical NPCs results in displacement of the mitotic cleavage plane with abnormal distribution of contractile components, we assessed Anillin distribution in *Lis1-MADM* neocortices compared with those of *WT-MADM* at E14.5. In the *WT-MADM* neocortex, Anillin was accumulated at the midzone during metaphase-to-anaphase (**Fig. 1 A-a,b**) and was enriched by forming a ‘U’ shape (basal-to-apical ingression) at the midbody of NPCs, consistent with previous observations of normal NPC cleavage in WT mice (Kosodo et al., 2008) (**Fig. 1 A-c,d**). In the *Lis1-MADM* neocortices, the tdTomato-positive WT NPCs (red, *Lis1^+/+^*) similarly displayed a normal distribution of Anillin at the apical cleavage furrow between the two daughter cells (**Fig. 1 C-e**). The tdTomato- and GFP-double-labeled *Lis1* heterozygous NPCs (yellow, *Lis1^ko/+^*) did not display an Anillin-rich ring associated with the midbody (**Fig. 1 C-f,g**). The *Lis1-MADM* (**Fig. 1 B-C**) neocortex displayed a profound decrease in GFP-positive *Lis1* homozygous KO apical NPCs located at the ventricular zone (green, *Lis1^ko/ko^*). These apical *Lis1^ko/ko^* NPCs were mostly found at prometaphase or metaphase and located at the ventricular surface with no obvious cell membrane-associated Anillin with dispersed patterns (**Fig. 1 C-h**), probably due to mitotic arrest after complete loss of LIS1 (Yingling et al., 2008). Abnormal distribution of Anillin in *Lis1* mutant NPCs (*Lis1^ko/+^*) implies that LIS1 is an important cell determinant for neocortical NPC cell cleavage and daughter cell separation during cytokinesis in a cell-intrinsic manner.

In apical NPCs of the neocortex, the highest expression of RhoA was detectable at the ventricular surface where NPC cleavage furrows were found (Gauthier-Fisher et al., 2009; Lian et al., 2019). Overexpression of dominant-negative forms of RhoA in mouse NPCs *in vitro* induced mislocalization of cleavage furrow with diffuse and dispersed contractile ring (Lian et al., 2019), suggesting that RhoA dictates cytokinetic progression and cleavage furrow specification. In the present study, cell membrane-bound forms of RhoA in apical NPCs were identified by fixing the embryonic brains with a 10% trichloroacetic acid (TCA) solution. In the control *WT-MADM* neocortex (*MADM-11^GT/TG^;Emx1-Cre^Cre/+^*), tdTomato illuminated the cytoplasm of dividing NPCs. At metaphase, RhoA was associated with the apical cell membrane of WT NPCs proximal to the ventricular surface (**Fig. 1 D-i**). At telophase, RhoA was enriched at the midbody located at the ventricular surface, indicating separation of the two daughter cells (**Fig. 1 D j,k,l**). However, in the *Lis1-MADM* neocortex (*MADM-11^GT/TG,Lis1^;Emx1-Cre^Cre/+^*), tdTomato- and GFP-double-labeled *Lis1* heterozygous NPCs (yellow, *Lis1^ko/+^*) displayed an unequal distribution of RhoA skewed to the cytoplasm of only one daughter cell (**Fig. 1 E-m,n**). The GFP-positive *Lis1* KO NPCs (green, *Lis1^ko/ko^*) were sparsely found at the prometaphase or metaphase with apical membrane-associated RhoA (**Fig. 1 E-o**). Together, mislocalization of Anillin and RhoA in *Lis1* mutant neocortical NPCs (*Lis1^ko/+^*) indicates that LIS1 may contribute to cleavage plane specification and positioning during late stages of apical NPC mitosis.

### Mislocalization of Anillin in apical NPCs from *Lis1* heterozygous neocortex

We next asked whether *Lis1* heterozygosity leads to changes in cytoarchitecture of the apical NPC niche at the ventricular surface of the neocortex. We deleted one copy of *Lis1* in neocortical NPCs by mating *Lis1* conditional knock-out (CKO) line with the *hGFAP-Cre* line (Zhuo et al., 2001). In control neocortex (*Lis1^hc/+^* without Cre, hc: hypomorphic conditional), NPCs undergoing vertical divisions (with a vertical cleavage plane angle) possessed basally located Anillin that gradually ingressed apically with the formation of the midbody, defining final location of cleavage furrow (basal-to-apical ingression) (**Fig. 2 A**), consistent with previous findings (Kosodo et al., 2008). NPCs undergoing horizontal or oblique divisions maintained a straight line of Anillin-rich immunoreactive signal in the junctional plate between two daughter cells (**Fig. 2 B**), suggesting normal cleavage furrow formation at the midzone between the two daughter cells. There were reduced numbers of NPCs undergoing vertical divisions (42%) in NPC-specific *Lis1* heterozygous mutant neocortex (*hGFAP-Cre; Lis1^hc /+^*) compared with those numbers (84%) in control neocortex (*Lis1^hc/+^*), consistent with our previous findings (Yingling et al., 2008). In addition, NPCs undergoing oblique and horizontal divisions in *Lis1* heterozygous mutant neocortex displayed a diffuse pattern of Anillin associated with similar enrichment at both the apical and basal cell membrane. Only one apically attached daughter cell retained concentrated Anillin-positive puncta at the ventricular surface (**Fig. 2 B-f,g,i**), suggesting that *Lis1* heterozygosity resulted in Anillin mislocalization during late mitosis of apical NPCs.

**Fig. 2.**
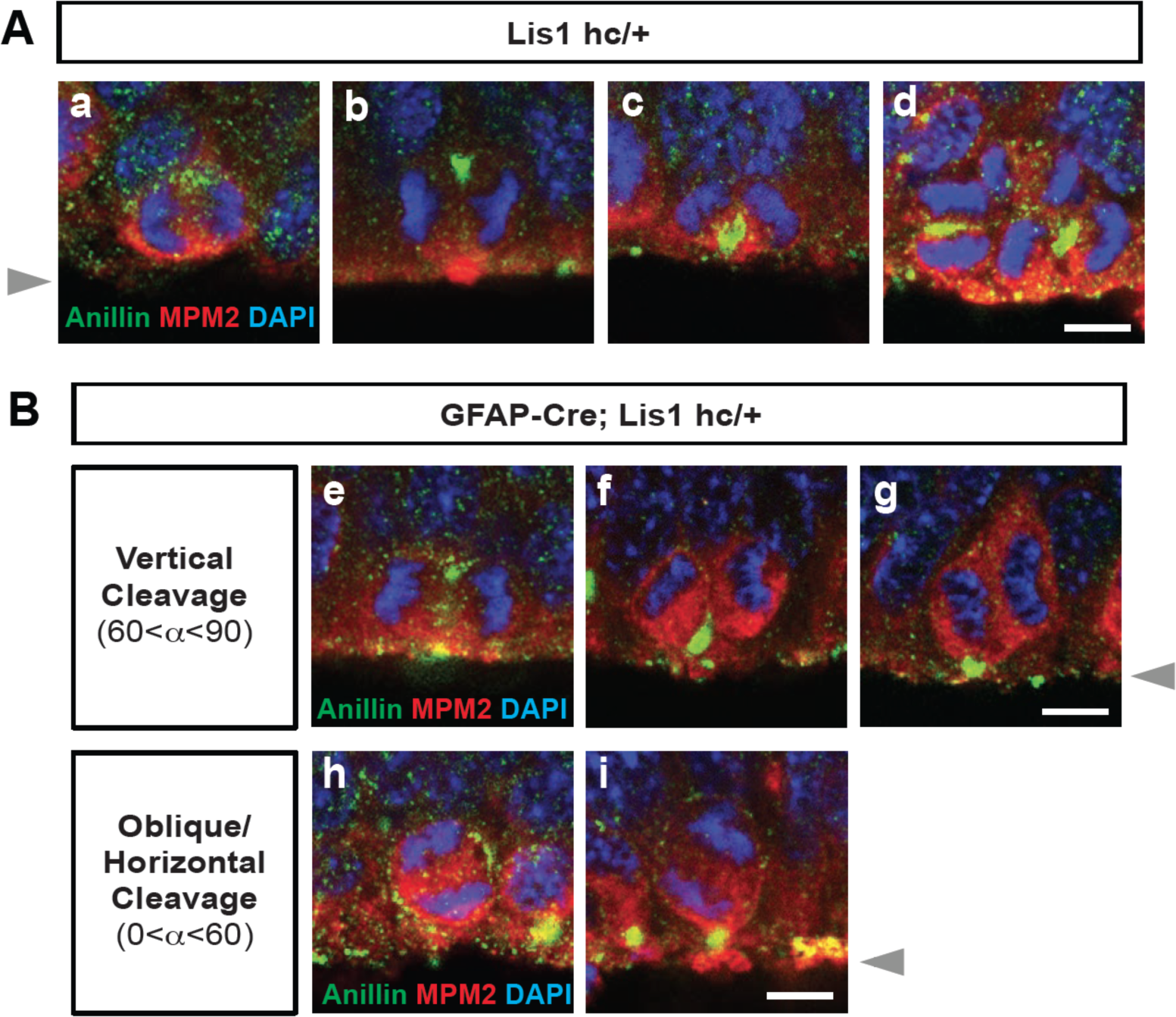
Localization of Anillin-ring in neocortical NPCs from controls and GFAP-Cre-induced *Lis1* mutant mouse embryos. **(A)** Normal Anillin-ring distribution during cytokinesis of *Lis1* control NPCs (*Lis1^hc/+^*). hc: hypomorphic conditional allele, **a.** Early anaphase, **b.** Mid-anaphase with a basally located Anillin-ring, **c.** ‘U’-like shape Anillin-ring at the cleavage furrow, **d.** Midzone-specific Anillin localization. **a.b.c.** Vertical cleavage plane (60°<α<90°), α: spindle angle compared to the ventricular surface, **d.** Oblique and horizontal cleavage plane (0°<α<60°). **(B)** Abnormal Anillin-ring localization during cytokinesis of *Lis1*-deficient mutant NPCs (*GFAP-Cre; Lis1^hc/+^*). **e.** Basal and apical Anillin-rings at the equator of NPCs, **f.g.** Moderately skewed Anillin-ring localization in only one daughter cell. h. The equator-associated Anillin-ring was detected with tilted spindle angle in *Lis1*-deficient mutant NPCs. **i.** *Lis1*-deficient mutant NPCs displayed horizontal cleavage plane with apical memrbrane-associated Anillin puncta. Grey arrowheads: ventricular surface (apical). Scale bars: 5 μm.

### An imbalance in symmetric versus asymmetric NPC divisions after dose-dependent reduction of LIS1

We hypothesized that *Lis1* mutant NPCs may display an imbalance in symmetric vs. asymmetric divisions due to defects in cleavage furrow patterning and cell membrane contractility. We previously reported that neocortical NPCs in *Lis1* mutants had reduced mitotic spindle angles because more cells displayed abnormal cleavage planes that were skewed to oblique and horizontal divisions (Yingling et al., 2008). Here, we analyzed neocortical NPC divisions by tracing the inheritance of atypical protein kinase C zeta (aPKC, PKCζ), an apical complex component, into each daughter cell. The cleavage furrow was identified by the location of Cadherin hole in the continuous line of N-Cadherin that indirectly indicates the contractile ring positioning (Marthiens and ffrench-Constant, 2009). The relative positioning of the mitotic spindle was defined by the midline between two chromosome sets. Thus, we determined that neocortical NPCs from *Lis1*-deficient mutants (*Lis1^hc/ko^*) more frequently displayed unequal inheritance of aPKCζ to the daughter cells (**Fig. 3 A-C**). In addition, apical NPCs in *Lis1^hc/ko^* neocortices possessing horizontal cleavage planes displayed less polarized aPKCζ along the apical-basal axis (**Fig. 3 B**), indicating mild alterations of apical polarity in *Lis1*-deficient neocortical NPCs. In *Lis1^hc/+^* control neocortices, 30.7% of NPCs displayed unequal distribution of aPKCζ into the cytoplasm of the two daughter cells (presumably asymmetric divisions). However, in *Lis1^hc/ko^* brains, 68.8% of NPCs displayed unequal distribution (**Fig. 3 C**). These results suggest that *Lis1* deficiency in neocortical NPCs results in abnormal cleavage furrow positioning that may induce unequal inheritance and distribution of cell fate determinants into two daughter cells that ultimately leads to an increase in asymmetric (presumably neurogenic) NPC divisions.

**Fig. 3.**
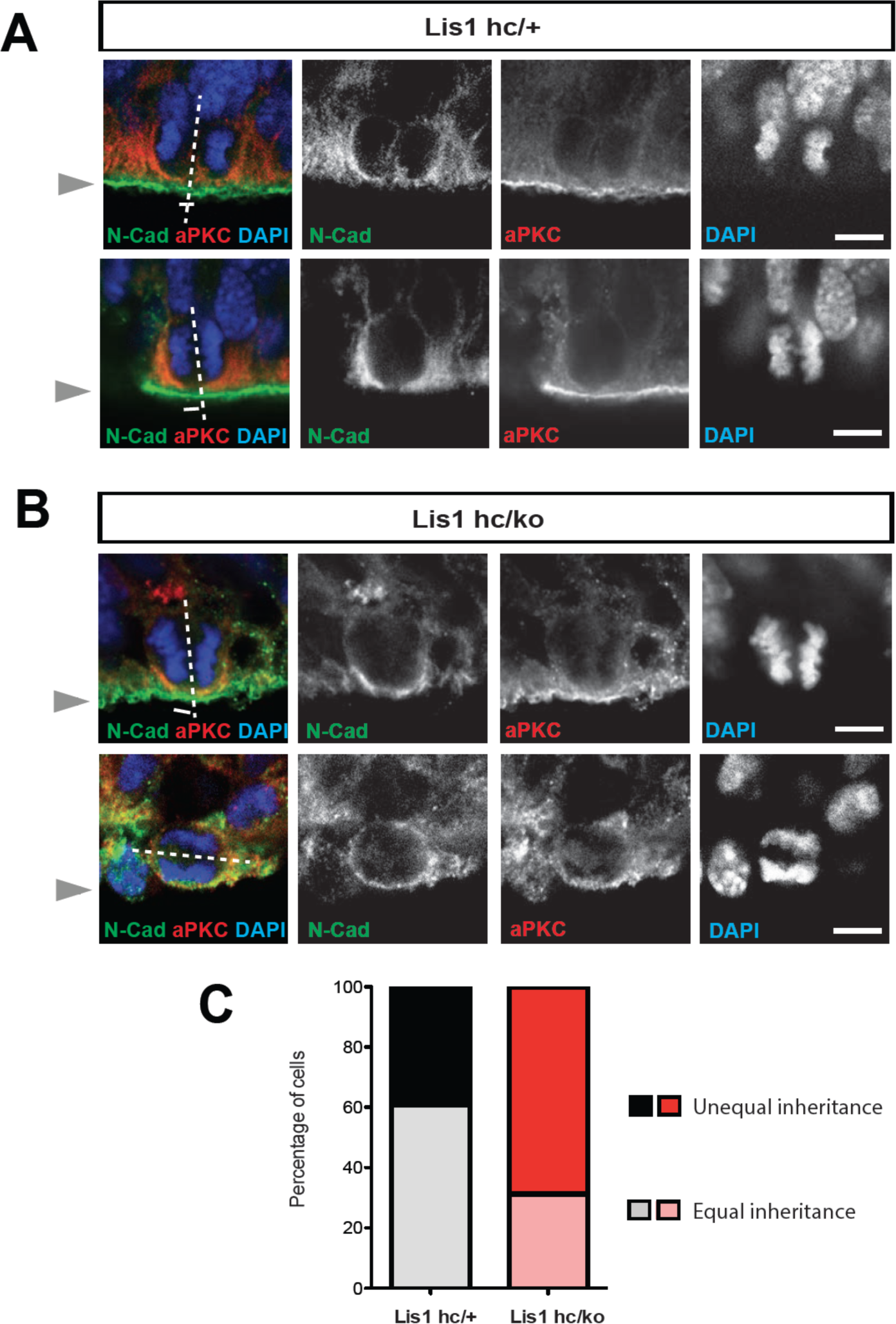
Alterations in symmetric and asymmetric divisions determined by equal vs. unequal inheritance of atypical PKC (aPKC, PKCz) and Cadherin holes. **(A)** Normal cytokinesis in neocortical NPCs from *Lis1* controls (*Lis1^hc/+^*). Upper panel: Control NPCs displayed vertical cleavage with equal inheritance of aPKC and Cadherin hole. Lower panel: Control NPCs displayed vertical cleavage with unequal inheritance of aPKC and Cadherin hole – presumably reflecting an asymmetric neurogenic division. **(B)** Upper panel: *Lis1*-deficient NPCs (*Lis1^hc/ko^*) displayed vertical cleavage with unequal inheritance of aPKC and Cadherin hole. Lower panel: *Lis1*-deficient NPCs displayed horizontal cleavage with unequal and very skewed inheritance of aPKC and Cadherin hole. This *Lis1*-deficient NPC retained less polarized aPKC/N-Cadherin protein distribution (less apical enrichment of aPKC and N-Cadherin) along the apical-basal axis. **(C)** Disturbances in the frequencies of equal vs. unequal inheritance of cell fate determinants in *Lis1*-deficient NPCs. Grey arrowheads: ventricular surface (apical). Scale bars: 5 μm.

### Perturbed daughter cell separation and an increase in binucleation during mitosis of ***Lis1* mutant MEF**

We also tested whether *Lis1* deficiency impairs cleavage plane positioning and completion of mitosis in MEFs using time-lapse microscopy. We visualized mitotic events of late stages of mitosis of MEFs *in vitro* in greater detail than we could do in the neocortex *in vivo*. We examined the frequency of normal daughter cell separation with complete abcission from both WT MEFs (*Lis1^+/+^*) and mutant *Lis1* compound heterozygous MEFs (*Lis1^hc/ko^*) expressing 35% of LIS1 compared to normal WT levels (Yingling et al., 2008). Sequential mitotic events were monitored by time-lapse live-cell imaging using mCherry-α-Tubulin, a fluorescently labeled MT marker, and histone 2B (H2B)-GFP, a chromosomal probe (Moon et al., 2014). A majority of WT MEFs displayed successful daughter cell separation (85.4%), while in *Lis1^hc/ko^* MEFs, this frequency was decreased (18.7%) (**Fig. 4 A**). By contrast, binucleated daughter cells was increased in *Lis1^hc/ko^* MEFs (40.1%) relative to WT MEFs (5.1%) (**Fig. 4 B**). To assess whether an increase in incomplete cell separation leads to changes in apoptotic cell death, we immunostained cleaved caspase-3. *Lis1^hc/ko^* MEFs had an increased number of cleaved caspase-3-immunoreactive cells compared with WT, suggesting that *Lis1* deficiency induces more frequent apoptosis of MEFs (**Fig. 4 C**). We also confirmed an increase in cell death (from 3- to 6-fold) from live-cell imaging of *Lis1^hc/ko^* MEFs.

**Fig. 4.**
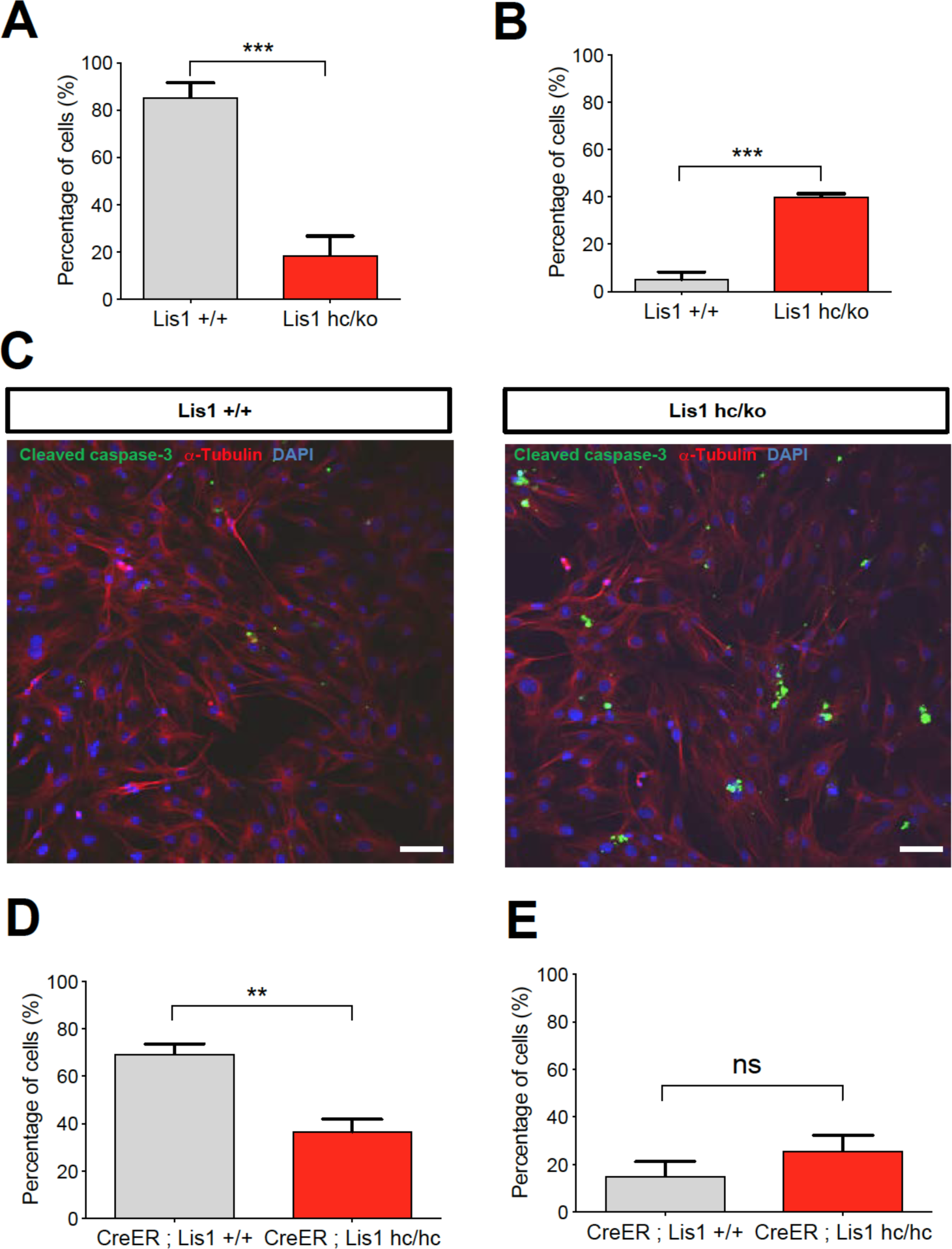
Loss of LIS1 in mouse embryonic fibroblasts (MEFs) impairs normal cytokinesis and leads to binucleation of the daughter cells. **(A)** Reduced number of *Lis1*-deficient mutant mouse embryonic fibroblasts (MEFs) (*Lis1^hc/ko^*) underwent normal cytokinesis with completion of daughter cell separation. *Lis1*-deficient mutant MEFs displayed frequent failure of cytokinesis compared with WT MEFs (*Lis1^+/+^*), analyzed by time-lapse live-cell imaging. **(B)** Increased binucleation by fused daughter cells from *Lis1*-deficient mutant MEFs (*Lis1^hc/ko^*) compared with WT MEFs. **(C)** Increased apoptotic cell death from *Lis1^hc/ko^* MEFs relative to WT MEFs (green: cleaved caspase-3, red: a-Tubulin, blue: DAPI). **(D)** Acutely induced *Lis1* conditional knock-out (CKO) mutant MEFs (*CreER^TM^; Lis1^hc/hc^ +* TM^12h^) also displayed less frequency of normal daughter cell separation compared with control MEFs (*CreER^TM^; Lis1^+/+^ +* TM^12h^). **(E)** An increased trend in binucleation was found in *Lis1* CKO mutant MEFs relative to control MEFs. However, it did not reach significant. Student’s *t*-test, \*\**p* <0.01, \*\*\**p* <0.001, ns: not significant, Scale bars: 50 μm.

To determine whether acute loss of *Lis1* also induces defective cell separation, we derived MEFs containing CKO alleles of *Lis1* and tamoxifen (TM)-inducible *CreER^TM^* line (Hayashi and McMahon, 2002) with the genotype *CreER^TM^; Lis1^hc/hc^*. We treated these MEFs with 4-hydroxy-TM for 12 hours and compared mitotic phenotypes with control MEFs (*CreER^TM^; Lis1^+/+^*). Similar to *Lis1^hc/ko^* mutant MEFs, acute deletion of *Lis1* in TM-treated *Lis1* CKO MEFs (*CreER^TM^; Lis1^hc/hc^* +TM^12 h^) induced a decrease in normal daughter cell separation (36.6%) compared with control levels (69.4%) (**Fig. 4 D**). Consistently, a trend of frequent daughter cell separation failure and higher percentage of binucleation (23.5%) than controls (*CreER^TM^; Lis1^+/+^* +TM^12 h^) (3.7%) was observed from live-cell imaging (**Fig. 4 E**) but this was not statistically significant.

### Cell shape oscillation and mitotic spindle rocking during mitosis of *Lis1* mutant MEFs

We analyzed movements of MT-enriched midbody and cleavage plane positioning throughout the entire mitotic cell cycle by live-cell imaging. Acute deletion of *Lis1* in MEFs (*CreER^TM^; Lis1^hc/hc^* +TM^12 h^) resulted in drastic cell shape oscillation (**Fig. 5 A, Movie S1**), similar to the phenotypes described in Anillin-depleted cells (Echard et al., 2004; Kechad et al., 2012; Piekny and Glotzer, 2008; Straight et al., 2003; Zhao and Fang, 2005). At the anaphase onset of *Lis1* mutant MEFs, the midbody was properly positioned in the central spindle zone. However, the position of the midbody was gradually destabilized and began vigorously oscillating between the two daughter cells. Surprisingly, the initially separated chromosome sets moved back and forth between two daughter cells and mitotic spindle rocking was prominent throughout this process. This phenotype was previously termed “hyper-contractility” in cells with depleted contractile ring components, and is caused by hyper-activity of the actomyosin during cytokinesis (Werner and Glotzer, 2008). Surprisingly, in one example seen in *Lis1* mutant MEFs (**Fig. 5 B, Movie S2**), one daughter cell inherited two-sets of chromosomes with binucleation while the other daughter cell underwent cell death processes.

**Fig. 5.**
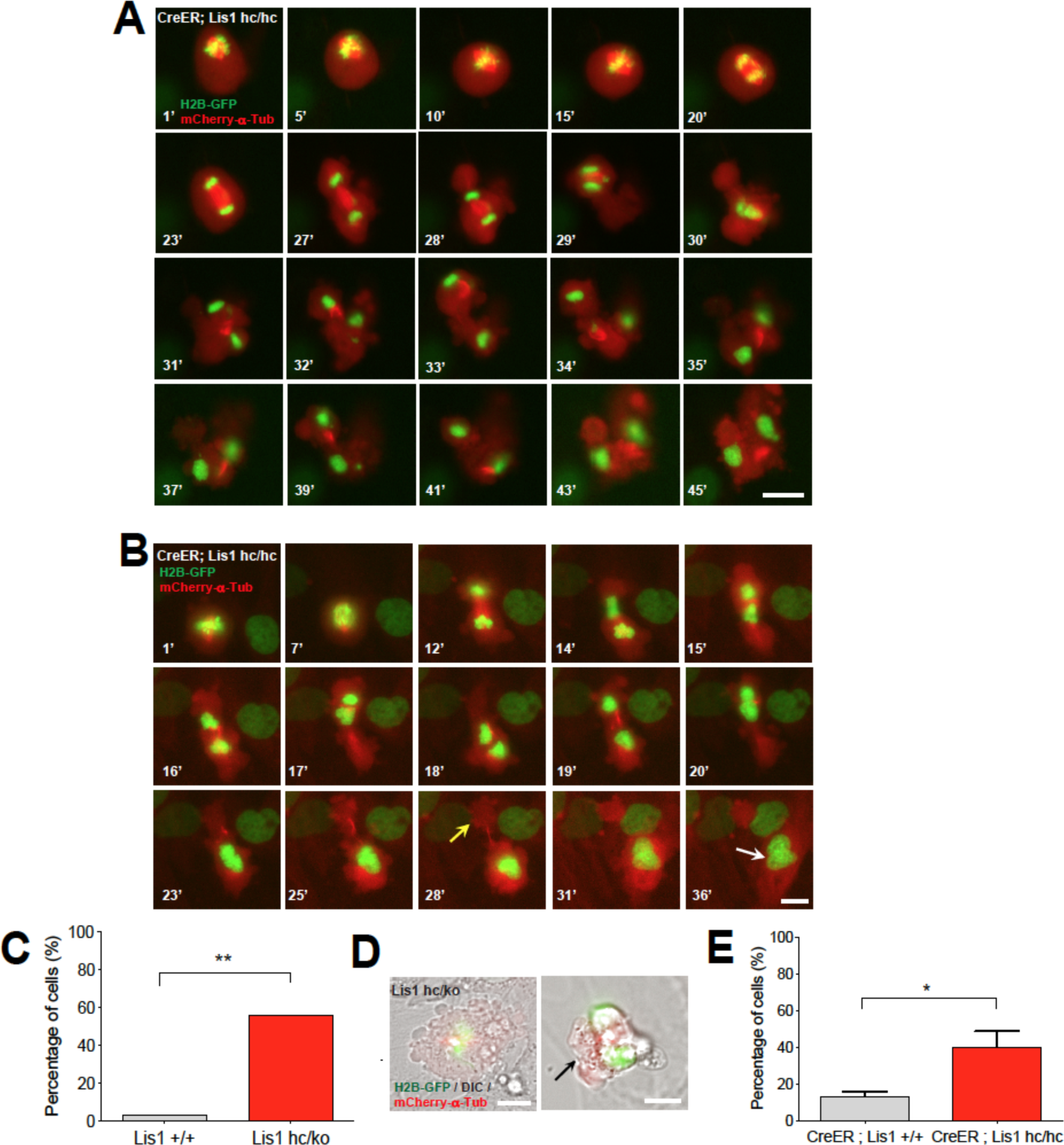
Loss of LIS1 results in cell shape oscillation and mitotic spindle rocking that leads to cytokinetic failure with aberrant cleavage plane positioning. **(A)** Acutely deleted *Lis1* mutant MEFs (*CreER^TM^; Lis1^hc/hc^ +* TM^12h^) displayed severe cleavage plane defects during cytokinesis. There was instability of the midbody (highly concentrated with red fluorescence: mCherry-α-Tubulin) accompanied by vigorous cell shape oscillation and mitotic spindle rocking (green: H2B-GFP). Scale bar: 20 μm. **(B)** The MEFs with acutely deleted *Lis1* displayed abnormal cell shape and chromosome oscillation between two daughter cells. Yellow arrow (28’): anucleated one daughter cell, White arrow (36’): binucleated and bi-lobed daughter cell. Scale bar: 10 μm. **(C)** A increase in the frequency of hyper-contractility phenotypes in *Lis1^hc/ko^* MEFs. **(D)** Single snapshots from the merged images with DIC (differential interference contrast) and fluorescence images (green: H2B-GFP, red: mCherry-α-Tubulin, grey: DIC) from time-lapse movies (black arrows: aberrant formation of huge membrane blebs). Scale bars: 10 μm. **(E)** Hyper-contractility quantification indicates that *Lis1* CKO mutant MEFs have increases in cell membrane blebbing phenotypes than control MEFs. Student’s *t*-test, \**p* <0.05,\*\**p* <0.01.

We next examined the incidence of the hyper-contractility phenotypes during MEF mitosis. WT MEFs rarely displayed oscillated cell membrane movements (3.6%). By contrast, more than half of the mitosis (56.2%) in *Lis1^hc/ko^* MEFs exhibited the hyper-contractility phenotypes (**Fig. 5 C**). The incidence of ectopic membrane bulges was increased in *Lis1* mutant MEFs which prompted us to monitor cell shape changes and cell membrane dynamics by merging phase-contrast images in live-cell imaging analysis. We found that *Lis1^hc/ko^* MEFs frequently showed cell membrane blebbing (**Fig. 5 D**) and the size of ectopic membrane blebs was much larger than those in WT MEFs. In accordance with these findings, the hyper-contractility phenotypes with cell membrane blebbing were more frequent in *Lis1* CKO MEFs (*CreER^TM^; Lis1^hc/hc^* +TM^12 h^) (40.5%) compared with control MEFs (*CreER^TM^; Lis1^+/+^* +TM^12 h^) (13.5%) (**Fig. 5 E**).

### Mislocalization of RhoA and contractile ring components in mitosis of *Lis1* mutant MEFs

We tested whether critical regulators of cytokinesis contribute to *Lis1*-deficiency-induced cell membrane hyper-contractility. Since RhoA is an important master GTPase regulating cytokinesis, we examined RhoA localization in *Lis1* mutant MEFs fixed with 10% TCA (Yonemura et al., 2004). In WT MEFs, the active forms of RhoA symmetrically accumulated at both sides of the equatorial cortex and were concentrated in the midbody at late telophase and during cytokinesis (**Fig. 6 A**). From anaphase to telophase, RhoA colocalized with the cell membrane-associated Anillin, a key component of the contractile ring that defines the cleavage furrow ingression site. By contrast, 60% of *Lis1^hc/ko^* MEFs displayed mislocalization of the cleavage furrow with the formation of large polar blebs in ectopic membrane sites (**Fig. 6 B**). An aberrant cleavage furrow was found at only one side of the cell equatorial cortex of *Lis1^hc/ko^* MEFs, resulting in asymmetric mislocalization of RhoA and Anillin (**Fig. 6 B**, upper panel). The binucleated *Lis1^hc/ko^* MEFs displayed protruding polar blebs in ectopic locations (**Fig. 6 B**, lower panel). Intriguingly, RhoA immunoreactive signal decorated the cell membrane of all of these polar blebs, suggesting that RhoA and actomyosin may be the primary signals to produce aberrant membrane bulges in *Lis1^hc/ko^* MEFs.

**Fig. 6.**
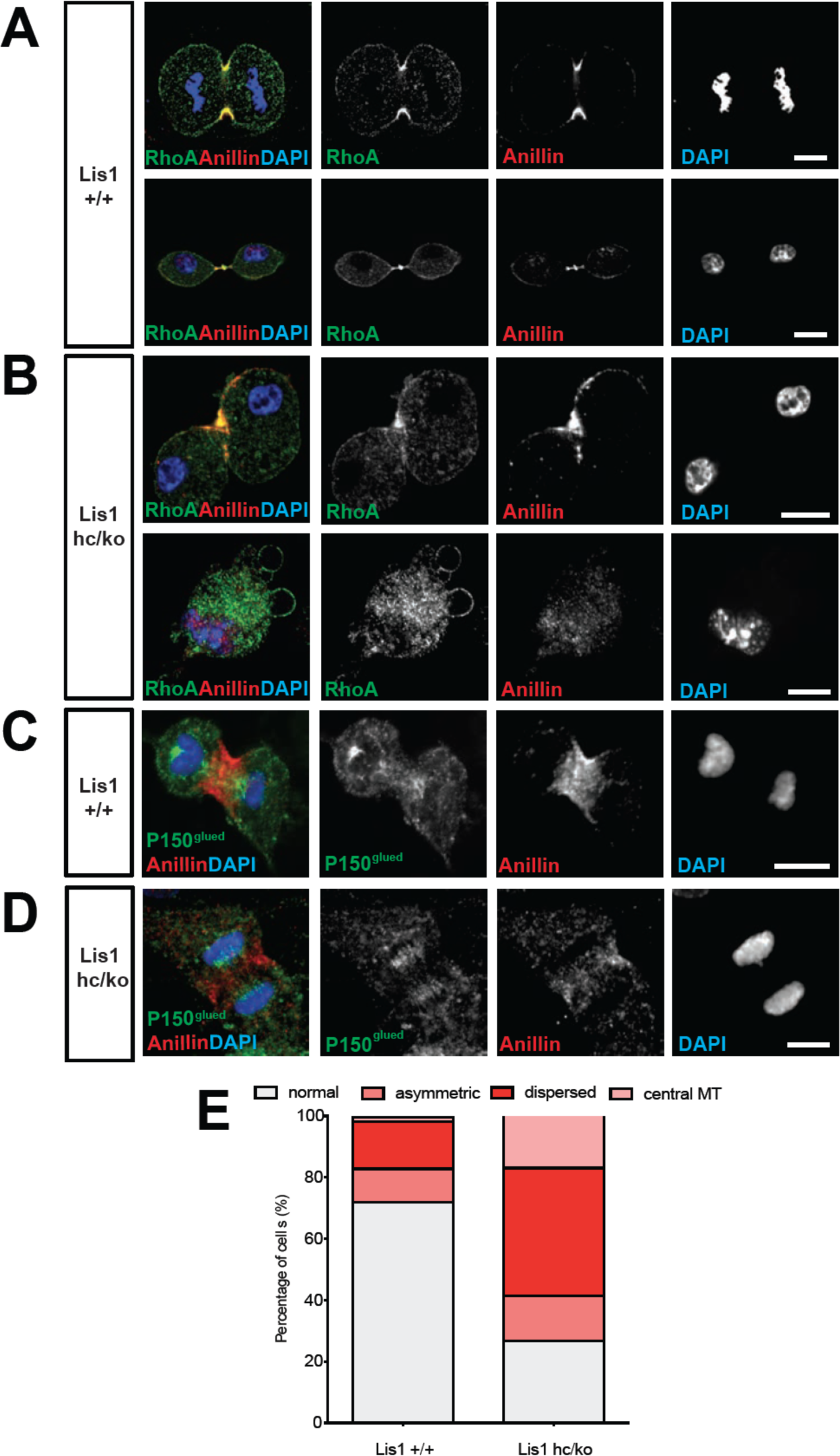
*Lis1* mutant MEFs display mislocalization of RhoA and Anillin, an important contractile ring component. **(A)** WT MEFs displayed normal cleavage furrow positioning labeled with TCA-fixed RhoA and Anillin co-staining at the equatorial cortex from early anaphase to telophase. **(B)** *Lis1^hc/ko^* MEFs displayed mislocalization of RhoA and Anillin. Upper panel: asymmetric cleavage furrow formation. Lower panel: binucleated cells with RhoA-positive aberrant membrane blebs. **(A,B)** (green: RhoA, red: Anillin, blue: DAPI). **(C)** WT MEFs displayed an Anillin-positive zone in the cell equator. The polar cortex-associated cortical P150^glued^ dynactin staining pattern was evident and it did not overlap with the Anillin-ring. **(D)** *Lis1^hc/ko^* MEFs exhibited reduced cortical P150^glued^ dynactin at the polar cortex and also have dispersed Anillin distribution at the equatorial cortex. **(C,D)** (green: P150^glued^ dynactin, red: Anillin, blue: DAPI). **(E)** Quantification of normal, assymetric, dispersed, and central MT-associated Anillin distribution in MEFs. Scale bars: 10 μm.

Since the LIS1-dynein-dynactin complex is associated with the cortical cell membrane (Faulkner et al., 2000; Moon et al., 2014), it is plausible that cortical dynactin, P150^glued^, may be less enriched in the cell membrane and cause Anillin mislocalization in *Lis1^hc/ko^* MEFs. Cortical P150^glued^ was incorporated within the polar cortex but it was excluded in the cleavage furrow where Anillin was associated with the equatorial cortex in WT MEFs (**Fig. 6 C**). However, reciprocal exclusion of polar P150^glued^ vs. equatorial Anillin was less obvious in *Lis1^hc/ko^* MEFs (**Fig. 6 D**) (Moon et al., 2014). From anaphase to telophase, the cells with cleavage furrow-associated Anillin were reduced in *Lis1^hc/ko^* MEFs (27.1%) relative to WT control *Lis1^+/+^* MEFs (72.3%) (**Fig. 6 E**, in grey).

### Mislocalization of F-actin and Myosin II in mitosis of *Lis1* mutant MEFs

A previous study demonstrated that the leading process-associated F-actin is misregulated during migration of post-mitotic neurons in *Lis1^ko/+^* mutants (Kholmanskikh et al., 2003). Actin dysfunction induced by loss of LIS1 may lead to cytokinetic defects and hyper-contractility seen in *Lis1^hc/ko^* MEFs. To test this possibility, we co-immunostained MEFs with phalloidin (F-actin zone) and dynein intermediate chain (DIC 74.1). WT MEFs had an F-actin zone located at the midbody that also colocalized with DIC (**Fig. 7 A**, upper panel). By contrast, *Lis1^hc/ko^* MEFs had wider F-actin cortical patches at the midzone that extended to the polar cortex in one daughter cell compared with WT MEFs. Central MT-associated F-actin fibers were also evident in *Lis1^hc/ko^* MEFs (**Fig. 7 A**, lower panel).

**Fig. 7.**
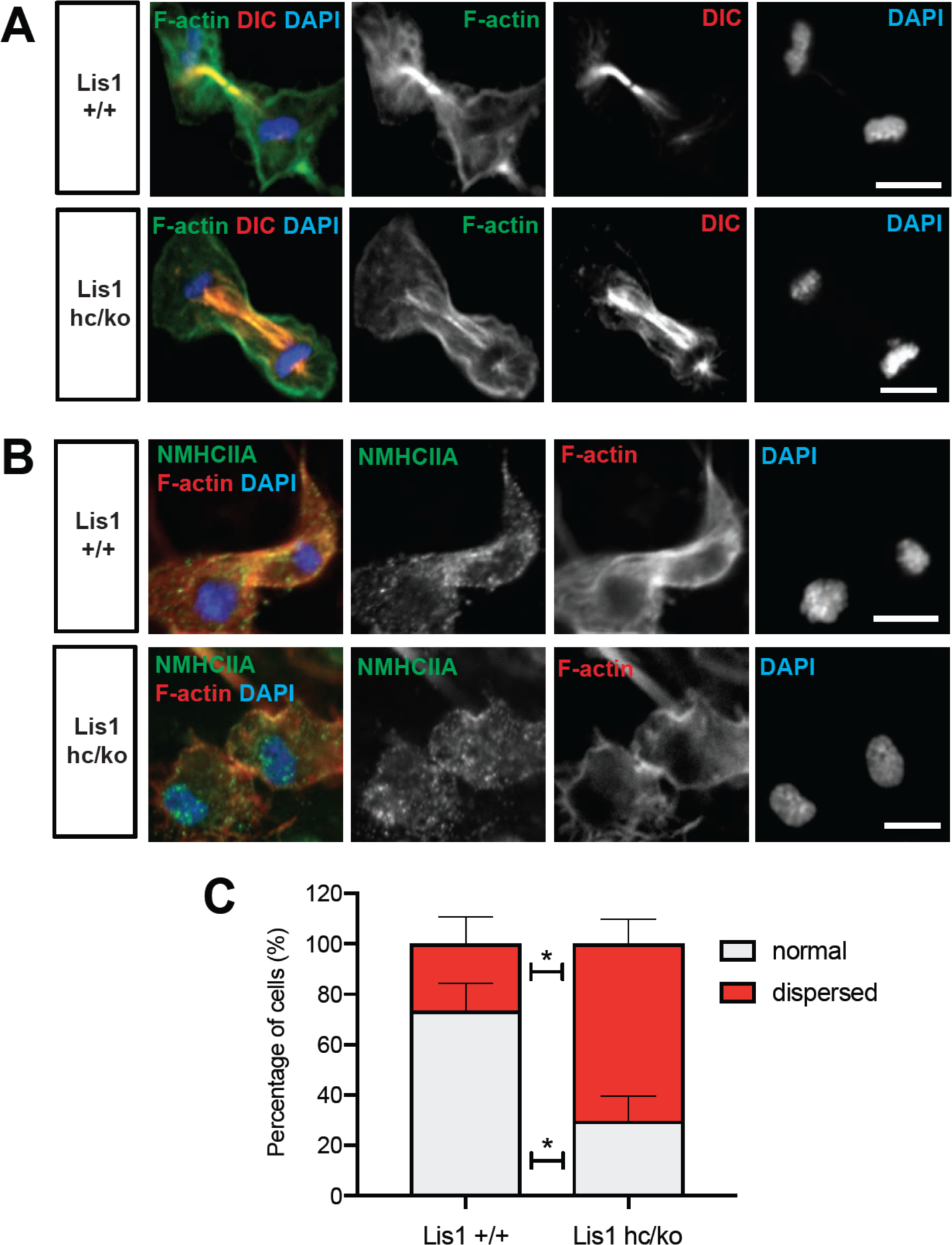
*Lis1* mutant MEFs show alterations in F-actin focus zone and Myosin II motor localization during late stages of mitosis. **(A)** From telophase to cytokinetic phases, WT MEFs maintain filamentous actin (F-actin) zone (phalloidin) at the midbody where the dynein complex (indicated by DIC, dynein intermediate chain) is accumulated. *Lis1^hc/ko^* MEFs exhibited defects to locate the dynein complex at midzone partly overlapped with widely dispersed F-actin zone. **(B)** WT MEFs with accumulation of non-muscle myosin heavy chain II-A (NMHCIIA) at the cell equator. Colocalization of staining of NMHCIIA and F-actin (phalloidin)-focusing zone (green: NMHCIIA, red: F-actin, blue: DAPI). *Lis1^hc/ko^* MEFs displayed dispersed and less-focused actomyosin at the cell equator. (green: NMHCIIA, red: F-actin, blue: DAPI). **(C)** Quantification of normal, assymetric, dispersed and central MT-associated NMHCIIA and F-actin distribution in MEFs at late stages of mitosis (at telophase or cytokinesis). Scale bars: 10 μm.

We next examined the localization of Myosin II in *Lis1^hc/ko^* MEFs relative to WT MEFs. Myosin II is the main molecular motor of the cortical contraction complex of cytokinesis, cooperatively working with the F-actin (Piekny et al., 2005). In WT MEFs, non-muscle myosin heavy chain II A (NMHCIIA) immunoreactive signal at the spindle midzone overlapped with the F-actin focus zone (**Fig. 7 B**, upper panel). By contrast, we found that MHCIIA distribution was diffused and dispersed at the spindle midzone in *Lis1^hc/ko^* MEFs (**Fig. 7 B**, lower panel). Quantitation of dispersed MHCIIA and F-actin distribution indicated that *Lis1^hc/ko^* MEFs had fewer cells with normal cleavage furrow-associated actomyosin patterns (*Lis1^+/+^*, 73.4% vs*. Lis1^hc/ko^*, 29.7% – in grey). Conversely, *Lis1^hc/ko^* MEFs showed an increase in cells with ectopic distribution of actomyosin (*Lis1^+/+^*, 26.6% vs. *Lis1^hc/ko^*, 70.3% – in red). (**Fig. 7 C**). These findings suggest that cortical constriction at the equatorial cortex was impaired in *Lis1^hc/ko^* MEFs, leading to mispositioning of the cleavage furrow by actomyosin dysfunction.

### Abnormal Myosin II movements during mitosis of *Lis1* mutant MEFs

We also monitored actomyosin dynamics by performing time-lapse live-cell imaging of non-muscle myosin regulatory light chain1 (MRLC1)-GFP in MEFs. MRLC1-GFP is a reliable molecular probe to illustrate the localization of the Myosin II (Beach et al., 2011; Miyauchi et al., 2006). We infected MEFs with MRLC1-GFP and H2B-tdTomato retroviruses to visualize Myosin II and chromosomes, respectively. In control MEFs (*CreER^TM^; Lis1^+/+^* +TM^24 h^), MRLC1 accumulated at the equatorial cortex and the contractile ring, then it constricted normally (**Fig. 8 A, Movie S3**). By contrast, Cre-inducible *Lis1* CKO mutant MEFs (*CreER^TM^; Lis1^hc/hc^* +TM^24 h^) displayed impairments in Myosin II movements (**Fig. 8 B, Movie S4**). Cleavage furrow ingression first occurred but regressed abnormally, with failure to restrict the cleavage furrow at the equatorial cortex. Cleavage furrow mispositioning occurred and resulted in chromosome missegregation. Multiple ectopic membrane bulges were found near the polar cortex in *Lis1* mutant MEFs, suggesting that aberrant cytoplasmic pushing forces may be generated by ectopically located Myosin II.

**Fig. 8.**
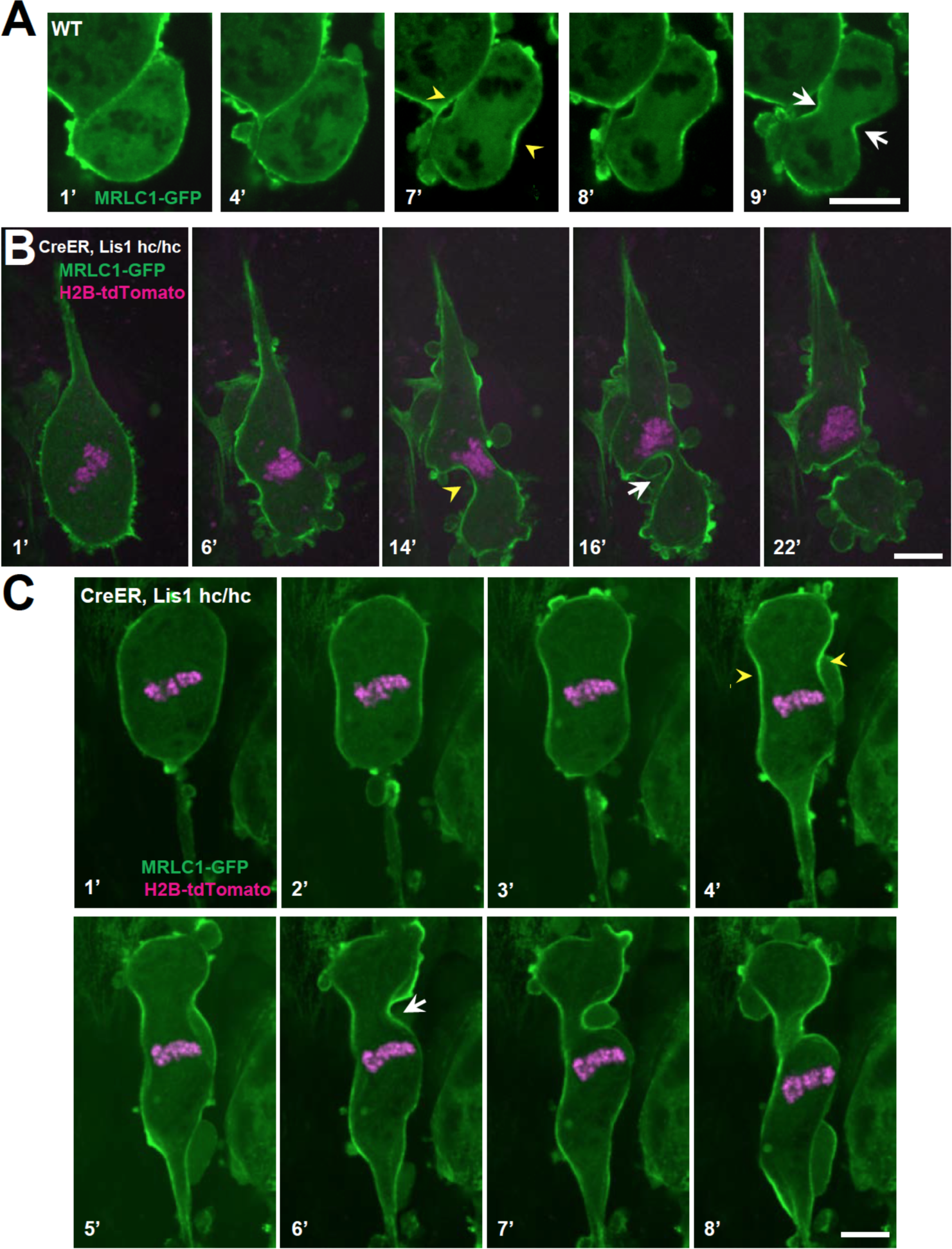
Myosin-II is abnormally distributed in *Lis1* mutant MEFs during late stages of mitosis. **(A)** WT MEFs displayed normal recruitment of the myosin regulatory light chain 1 (MRLC, green) of Myosin-II subunit at the equatorial cortex during cytokinesis. Dark black spots inside of the cells indicate chromosome sets. **(B)** *Lis1* mutant MEFs (*CreER^TM^; Lis1^hc/hc^ +* TM^24h^) infected with MRLC1-GFP (green) and H2B-tdTomato (magenta), displayed cytokinesis failure with an asymmetrically mis-positioned cleavage furrow. **(C)** *Lis1* mutant MEFs clearly displayed uncoupling between chromosome separation (usually before cytokinesis onset) and cleavage furrow ingression. *Lis1* mutant MEFs (4’) still maintained unsegregated tetraploid chromosomes to initiate furrow contraction from the actomyosin ring. **(A-C)** Yellow arrowheads: initial accumulation of MRLC1, White arrows: final cleavage furrow positioning. Scale bars: 5 μm.

In the severely affected cells, *Lis1* mutant MEFs displayed an uncoupling between chromosome segregation and cytokinesis (**Fig. 8 C, Movie S5**). In normal cytokinesis, chromosome sets first separated into the two daughters, followed by equatorial constriction to initiate cytokinesis. However, this sequential progression of cytokinesis was disrupted in *Lis1* mutant MEFs. Although chromosome sets did not segregate precisely into two daughter cells before the anaphase onset of mitotic spindle elongation, actomyosin-mediated hyper-contractility of the cell membrane triggered cytokinesis progression, producing binucleated cells.

### Failure to maintain contractile ring components at the equatorial cortex in *Lis1* **mutant MEFs**

To test whether the actomyosin dysfunction in *Lis1* mutant MEFs exert deleterious effects on the recruitment of contractile components to the equatorial cortex, we performed time-lapse live-cell imaging of Septin6 (SEPT6)-GFP (Gilden et al., 2012). Like Anillin, the Septin complex is the key component of the contractile ring during cell cleavage. In control MEFs, SEPT6 was enriched at the equatorial cortex and stayed at the cleavage furrow ingression site (**Fig. 9 A, Movie S6**). However, after acute *Lis1* deletion in *Lis1* CKO mutant MEFs (*CreER^TM^; Lis1^hc/hc^* +TM^24 h^), SEPT6 was initially observed at the equatorial cortex but then it regressed, and weak SEPT6 accumulation was detected at the cell equator accompanied by cortical deformation and chromosome oscillation/rocking (chromosomes were identified as darker spots in the SEPT6-GFP background) (**Fig. 9 B, Movie S7**). Further live-cell imaging of SEPT6-GFP infected MEFs showed that the frequency of cytokinetic failure with concomitant SEPT6 mislocalization was increased by 4.7-fold in *Lis1* mutant MEFs (66.7% in *Lis1^hc/ko^*) compared with WT controls (14.3% in *Lis1^+/+^*) (**Fig. 9 C**). Intercellular cytokinetic bridges and binucleation events resulted from incomplete cytokinesis were observed only from *Lis1*-deficient mutant MEF, not like WT MEFs. Together, these results indicate that the contractile ring components were not strictly retained at the equatorial cortex, similar to the defective localization of actomyosin, resulting in hyper-contractility and cytokinesis failure in *Lis1* mutant MEFs.

**Fig. 9.**
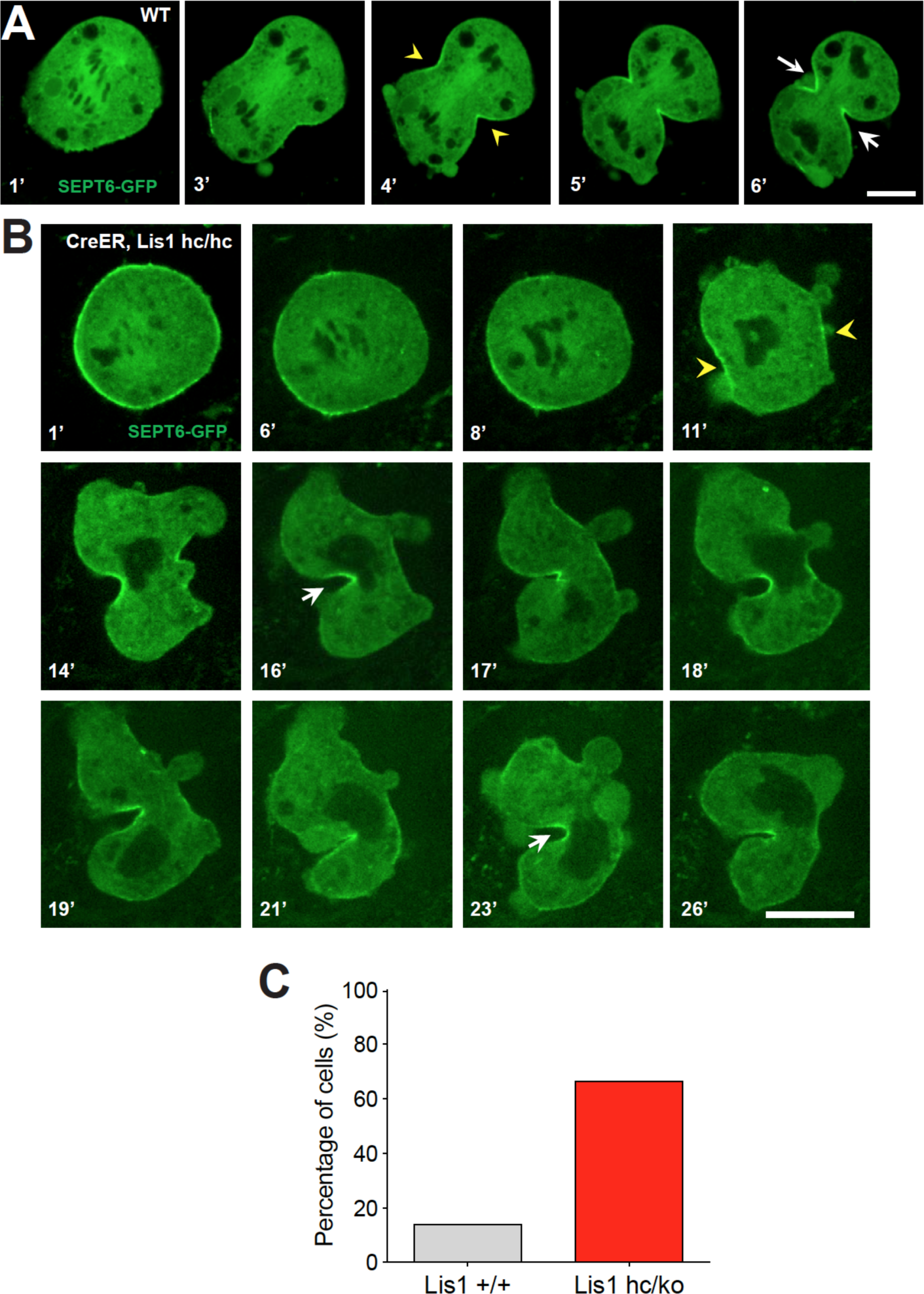
The contractile ring component SEPT6 is not properly maintained at the equatorial cortex in *Lis1* mutant MEFs. **(A)** WT MEFs displayed normal recruitment of Septin 6 (SEPT6-GFP, in green), an actomyosin-fiber crosslinking protein composed of contractile ring, to the equatorial cortex. **(B)** *Lis1* mutant MEFs (*CreER^TM^; Lis1^hc/hc^ +* TM^24h^) displayed vigorous cell shape oscillation with spindle/chromosome rocking. *Lis1* mutant MEFs displayed chromosome missegregation and failure to maintain proper contractile ring contraction sites. Dark black spots inside of the cells indicate the chromosome sets. **(A,B)** Yellow arrowheads: initial accumulation of SEPT6, White arrows: final cleavage furrow positioning). **(C)** Quantification of cytokinetic failure with abnormal asymmetric SEPT6-GFP distribution along the cell equantor indicates that binucleation events and incomplete cytokinesis occur more frequently in *Lis1*-deficient mutant MEFs than WT control MEFs. Scale bars: 10 μm.

### Modulation of RhoA hyper-activity in WT and *Lis1* mutant MEFs

If RhoA was hyper-activated in *Lis1* mutant MEFs, the cells expressing a constitutively active form of RhoA (CA-RhoA) might mimic cytokinesis defects found in the *Lis1* mutant MEFs. To test this possibility, WT MEFs were infected with retrovirus encoding human CA-RhoA (CA-RhoA)-GFP fusion protein. As expected, CA-RhoA-GFP-infected WT MEFs (*Lis1^+/+^* + CA-RhoA-GFP) displayed an increased frequency of cytokinesis failure with mislocalization of Anillin, similar to *Lis1^hc/ko^* MEFs (**Fig. 10 A**). WT MEFs infected with GFP control retrovirus (*Lis1^+/+^* + GFP) displayed normal cytokinesis (**Fig. 10 B**)

**Fig. 10.**
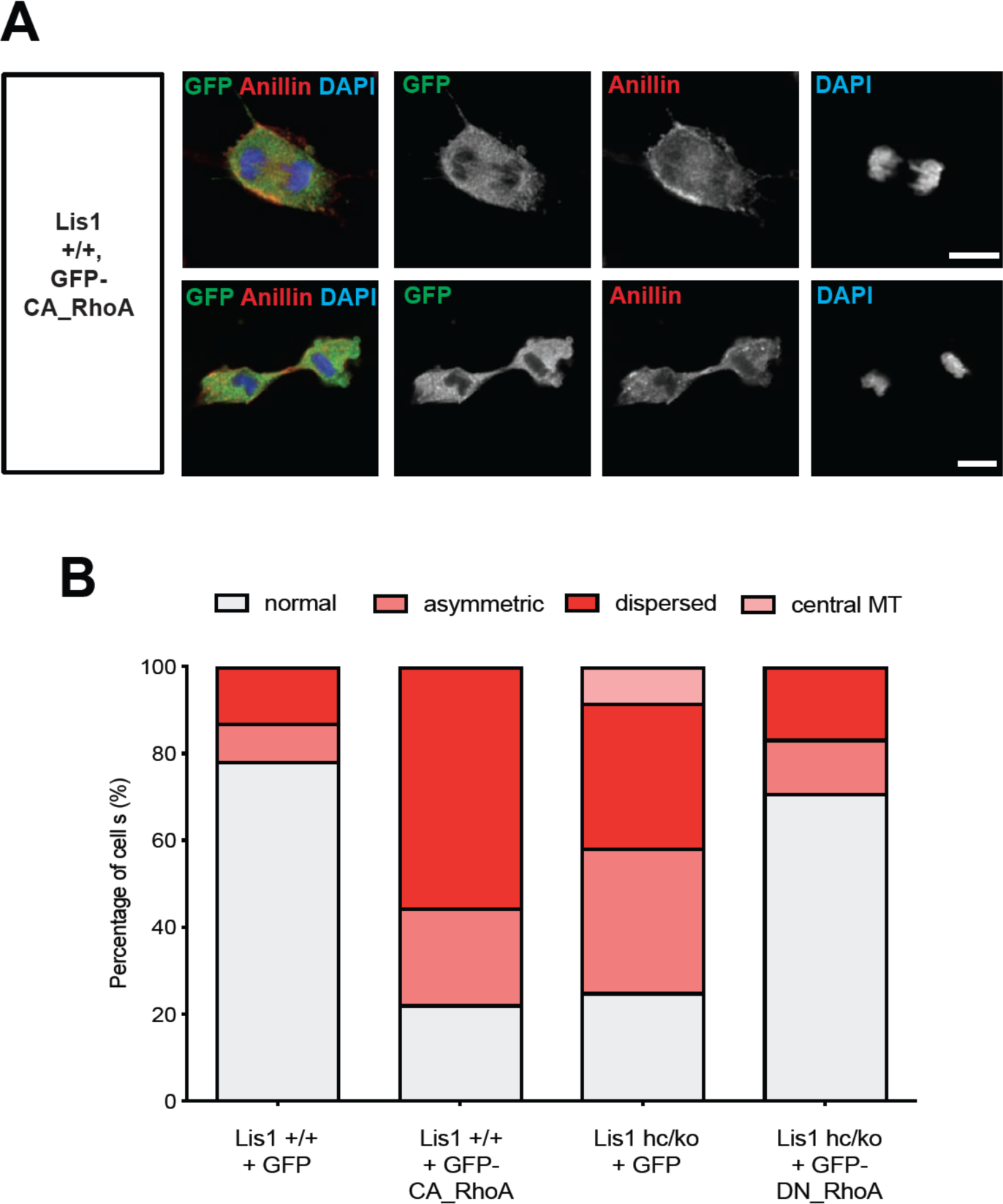
Overexpression of a constitutive active form of RhoA recapitulates cytokinetic failure similar to those are seen *Lis1* mutant MEFs and a dominant negative form or RhoA rescues hyper-activity of RhoA observed in *Lis1* mutant MEFs. **(A)** WT MEFs overexpressing a constitutive active form of RhoA (CA-RhoA) displayed asymmetric Anillin distribution at the cell equator at anaphase. Anillin recruitment to the midbody was not evident during cytokinesis of WT MEFs overexpressing CA-RhoA. **(B)** Quantification of Anillin distribution phenotypes in the late phases of mitoses of MEFs infected with GFP-fusion protein-expressing retroviruses. WT MEFs expressing CA-RhoA displayed abnormal Anillin distribution during cytokinesis consistent with cytokinesis defects seen in *Lis1^hc/ko^* MEFs. The *Lis1^hc/ko^* MEFs expressing a dominant negative form of RhoA (DN-RhoA) displayed partially rescued abnormal distribution Anillin, showing cytokinesis phenotypes similar to WT control MEFs. Scale bars: 10 μm.

Since *Lis1^hc/ko^* MEFs displayed the mitotic phenotypes resembling RhoA hyper-activity, we hypothesized that inhibition of RhoA by overexpression of a dominant negative form of RhoA (DN-RhoA) may ameliorate cytokinesis defects in *Lis1^hc/ko^* MEFs. *Lis1^hc/ko^* MEFs infected with retrovirus encoding a human DN-RhoA (DN-RhoA)-GFP fusion protein (*Lis1^hc/ko^* + DN-RhoA-GFP) displayed a decreased frequency of cytokinesis defects compared with control MEFs (*Lis1^hc/ko^* + GFP) (**Fig. 10 B**). These results imply that *Lis1^hc/ko^* MEFs displayed hyper-activation of RhoA and inhibition of RhoA that enabled the rescue of cytokinesis defects seen in *Lis1^hc/ko^* MEFs.

## DISCUSSION

Here we provide new evidence that LIS1 plays important roles in regulating cleavage plane positioning at late stages of mitosis during early mouse development. Normal levels of LIS1 expression are required for restricting cleavage furrow located at the equatorial cortex by inhibiting cell membrane hyper-contractility. It has been well understood that LIS1 is essential for mitotic spindle orientation by controlling MTs and dynein complex (Faulkner et al., 2000; Yingling et al., 2008; Moon et al., 2014). In addition, diverse cellular functions of LIS1 have been extensively studied in the migrating post-mitotic neurons (Gambello et al., 2003; Hirotsune et al., 1998; Sasaki et al., 2000; Tsai et al., 2005; Youn et al., 2009; Hippenmeyer et al., 2010). However, the detailed mitotic functions of LIS1 in cleavage plane specification during cytokinesis of daughter cell separation and its roles in actomyosin-mediated cell membrane contractility regulation have not been clearly elucidated.

In interphase cells and post-mitotic neurons, LIS1 has been known to modulate GTPase-mediated F-actin dynamics. *Lis1* heterozygous (*Lis1^ko/+^*) post-mitotic neurons displayed impairments in F-actin associated with the leading processes and dysregulated GTPase activities of RhoA-Rac1-Cdc42. Similarly, *Lis1^ko/+^* MEFs also exhibited abnormal hyper-activity of RhoA GTPase (Kholmanskikh et al., 2003). A recent *Lis1* knockdown fibroblast study unraveled F-actin dysfunction and impaired focal adhesion behaviors during active traction-dependent migration (Jheng et al., 2018). Together, these previous findings indicate that LIS1 may regulate not only MTs but F-actin in migrating phases of many cell types. However, whether LIS1 contributes to F-actin dynamics regulation during any of mitotic phases has not been investigated. Based on the aberrant cytokinetic phenotypes found in *Lis1* NPCs and MEFs described in this study, we demonstrated that loss of LIS1 results in hyper-active RhoA to transmit an abnormal signal to F-actin and myosin that generates hyper-contractility forces ectopically at the polar cortex, which is not a normal equatorial cortex location of cleavage furrow ingression.

Multiple mouse studies have indicated that RhoA GTPase regulates the ventricular surface integrity of the apical NPC niche, through controlling F-actin at the adherens junction (Cappello et al., 2012; Herzog et al., 2011; Katayama et al., 2011;). During chicken neural tube development, changes in RhoA activity and overexpression of a dominant negative form of RhoA similarly altered mitotic spindle orientation of apical NPCs (Roszko et al., 2006). However, it is unknown whether RhoA-mediated actomyosin regulation contributes to the later mitotic events when the segregation of cell fate determinants occurs. In the present study, we demonstrated that the LIS1-RhoA-actomyosin pathway also regulates apical NPC cleavage plane by recruiting the mediators of cytokinesis to the midbody at the ventricular surface. We found that *Lis1* mutant NPCs displayed mislocalization of RhoA and Anillin and this may impact the segregation of cell fate determinants by inducing an imbalance of symmetric vs. asymmetric divisions.

Interestingly, the exact cellular mechanisms that regulate cleavage plane positioning and furrow ingression during mitosis of apical NPCs remain mostly elusive. It appears that Anillin-ring constriction from the basal to apical side determines the cleavage furrow ingression site in mouse apical NPCs (Kosodo et al., 2008). It has been reported that Citron kinase and RhoA colocalize at the apical membrane of NPCs in the early rodent neocortices (Di Cunto et al., 2000; Sarkisian et al., 2002). Citron kinase mutations alter mitotic spindles and RhoA-mediated neurogenic cytokinesis of apical NPCs. These NPC phenotypes are associated with primary microcephaly and microlissencephaly in humans (Basit et al., 2016; Li et al., 2016; Shaheen et al., 2016; Harding et al., 2017; Gain and Di Cunto 2017). Through MT regulation and modulation of F-actin polymerization, three members of Kinesin family, Kif20a, Kif20b, and Kif4 are involved in neocortical NPC cytokinesis and mutations of these genes result in reduced cortical size with microcephaly (Janisch et al., 2013; Moawia et al., 2108; Geng et al., 2018). Together, these previous studies suggest that cooperation of MT and F-actin is the key process regulating NPC mitosis and cytokinesis by precisely segregating cell determinants along the cell cleavage axis.

In the present study, we demonstrated that LIS1 is an important mediator of signaling cascades of RhoA-actomyosin-contractile ring components during mitosis of apical NPCs, facilitating MDAM-mediated intrinsic pathway in the neocortex. To delineate mitotic behaviors affected by LIS1 deficiency, by performing live-cell imaging of MEFs, we discovered that *Lis1* mutant MEFs displayed abnormal phenotypes at the late stages of mitosis, cytokinesis, with increased incidence of binucleation and cell death resulted from the failure in cell separation. Cell shape oscillations and chromosome rocking along the mitotic spindles were also evident in *Lis1* mutant MEFs, surprisingly similar to those of Anillin-depleted cells (Echard et al., 2004; Kechad et al., 2012; Piekny and Glotzer, 2008; Straight et al., 2003; Zhao and Fang, 2005). Together, these results suggest that *Lis1* deficiency in MEFs induces cleavage furrow mispositioning and failure in the recruitment of contractile ring components to the equatorial cortex. Intriguingly, cell shape oscillation and cytokinesis defects were also previously reported in nocodazole-treated cells (Canman et al., 2003), suggesting that misregulation of either MT or the MT-actomyosin interface at the cell membrane may induce hyper-contractility leading to abnormal membrane contraction in *Lis1* mutant MEFs. *Lis1*-dependent cell membrane contractility may be mediated indirectly by the dyregulation of astral MTs (Yingling et al., 2008; Moon et al., 2014) reaching the cortical F-actin patch located in the cell membrane.

We aimed to identify the molecular mechanisms underlying *Lis1*-dependent cell membrane contractility, therefore, we examined distribution of the other proteins defining cleavage furrow including RhoA and its downstream effectors, like contractile ring components and actomyosin. We found that RhoA and Anillin were mislocalized in *Lis1* mutant MEFs. For instance, RhoA was found in ectopically localized large polar blebs in *Lis1* mutant MEFs. The Myosin II-positive zone and F-actin focus zone were dispersed at the equatorial cortex or aberrantly located at the polar blebs in *Lis1* mutant MEFs. By tracing MRLC1 and SEPT6, we confirmed that LIS1 defines cleavage plane positioning by dual regulation of actomyosin and contractile ring. Finally, RhoA hyper-activation in WT MEFs phenocopied the cytokinesis defects seen in *Lis1* mutant MEFs, and inhibition of RhoA in *Lis1* mutant MEFs decreased the cytokinesis defects observed in these MEFs. Thus, these results suggest that LIS1 exerts its function to restrict the cleavage furrow at the equatorial cortex to finely modulate RhoA-Anillin-actomyosin signaling cascades.

These findings also may explain our recent studies with induced pluripotent stem cells (iPSCs) of Miller Dieker syndrome, a severe form of lissencephaly with haploinsufficiency of *LIS1* as well as about 20 other genes. We demonstrated a prolongation of mitosis of outer radial glial (oRG) progenitors but not ventricular zone radial glial (vRG) progenitors (Bershteyn et al. 2017). The specificity of mitotic effects in the oRGs and lack of its effects in the vRGs may be the result of different dose dependencies of LIS1 in mouse vs. human, such that LIS1 deficiency results in more prolonged mitotic time specifically in oRGs than vRGs. Intriguingly, oRGs exhibit rapid somal translocation toward the cortical plate right before cytokinesis (Hansen et al., 2010). These abrupt movements suggest dynamic cytoskeletal rearrangements within mitotic oRGs. Since oRGs highly express a specialized transcriptional factor, HOPX (Pollen et al., 2015) on top of the pan-RG markers PAX6 and SOX2, it would be crucial to investigate the orchestrated crosstalk between HOPX-dependent transcriptional/epigenetic network and actomyosin-mediated cytokinesis in oRGs. In the future, it would be interesting to determine whether other lissencephaly or brain malformation-associated genes modulate RhoA-Anillin-actomyosin-pathways, and to test whether these genes, like LIS1, also possess dual roles in actomyosin and MT regulation important for apical NPCs by controlling key cellular processes not only mitosis but cytokinesis, a critical final step of cell determinant segregation.

## MATERIALS AND METHODS

### Animals

The mouse lines used in the present study were previously described; *CreER^TM^* (Hayashi and McMahon, 2002), *Lis1^hc/+^, ^hc/hc^* and *Lis1^hc/ko^* (Yingling et al., 2008; Moon et al., 2014), *hGFAP-Cre* (Zhuo et al., 2001), *MADM-11^GT/TG,Lis1^;Emx1-Cre^Cre/+^* (Hippenmeyer et al., 2010).

### Cell culture and Immunocytochemistry

Primary MEFs cell culture was performed as previously described (Moon et al., 2014). Cre recombinase in *CreER^TM^*-MEFs was induced by administration of 4-hydroxy tamoxifen (TM) (Sigma, 100 nM) for 12 hours. DMSO and blebbistatin (Calbiochem, 100 μM) were used to treat MEFs for 30 minutes. To identify RhoA in the cortex and cleavage furrow, MEFs were fixed with freshly made 10% trichloroacetic acid (TCA) for 10 minutes (Yonemura et al., 2004). Immunocytochemistry staining of cortical P150^glued^, LIS1, and NDE1 was performed in 100% methanol-fixed samples as previously described (Moon et al., 2014). The antibodies used in this study were listed in Table S1.

### Retrovirus infection, epifluorescence and spinning-disk confocal time-lapse live-cell imaging

Time-lapse live-cell imaging of MEFs infected with retroviruses encoding H2B-GFP, mCherry-α-Tubulin was described previously (Moon et al., 2014). To induce Cre-mediated *Lis1* CKO allele deletion, the MEFs derived from *CreER^TM^; Lis1^+/+^* and *CreER^TM^; Lis1^hc/hc^* animals were treated with 4-hydroxy-tamoxifen (TM) (Sigma, 100 nM) for 12 hours or 24 hours. To visualize Myosin II movements during cytokinesis, primary MEFs were co-infected with a mixture of MRLC1-GFP and H2B-tdTomato retroviruses with 4 µg/mL polybrene. Lasers with 488 nm and 561 nm emission were used for GFP and tdTomato imaging. SEPT6-GFP was also traced by a Nikon Ti spinning-disk confocal microscope with a 488 nm emission laser. The sequential 2 µm interval 5∼7 Z-stack images were captured every 30 seconds. The best Z confocal plane images showing the cleavage furrow were made as time-lapse movies. The plasmid sources of retrovirus construction were described in Table S1.

### Immunohistochemistry

Mouse fetal brains were isolated from E14.5 embryos (described in Yingling et al., 2008) followed by fixation with 4% paraformaldehyde (PFA) or 10% TCA (RhoA staining). On the next day, the brains were transferred to 30% sucrose and embedded in O.C.T. compound to make cryomolds. 18 μm-thick cryostat sections of neocortex were used for immunohistochemistry. In *MADM-11* animals, we identified *Lis1* genotype based on GFP or tdT immunoreactivity. Cleavage furrow markers such as Anillin and RhoA were immunostained in the far-red Alexa Fluor 647 channel to separate the fluorescence signal from GFP and tdT. Anillin- or RhoA-immunoreactive signals were pseudo-colored in green for representation purpose. The antibodies used in this study were listed in Table S1.

## ACKNOWLEDGEMENTS

We would like to thank Dr. Arshad Desai and Dr. Karen Oegema in UCSD for providing initial key insights into the novel phenotypes seen in *Lis1* mutant MEFs, Dr. Torsten Wittmann, Hayley Pemble, Kurt Thorn at Nikon imaging center in UCSF for access to microscopes for time-lapse live-cell imaging, Dr. Matthew Krummel in UCSF for Septin6-GFP plasmid, Dr. Kozo Kaibuchi in Nagoya University for RhoA plasmids and insightful advice, and Dr. Giles Plant in Stanford University for access to Nikon imaging analysis.

