## Supplementary Table S1 for "LIS1 determines cleavage plane positioning by regulating actomyosin-mediated cell membrane contractility"

### Table S1. List of antibodies used for immunostaining and retrovirus plasmids

- **Primary antibodies**

| Antibody name | Species | Vendor | Dilution |
|---|---|---|---|
| Cleaved caspase3 | rabbit | Cell Signaling | 1:500 |
| $\alpha$-Tubulin | rat | AbD Serotec | 1:1,000 |
| RhoA | mouse | Santa Cruz | 1:500 |
| Anillin | rabbit | Santa Cruz | 1:250 |
| P150$^{glued}$ | mouse | BD bioscience | 1:200 |
| DIC 74.1 | mouse | Millipore | 1:100 |
| NMHCIIA | rabbit | Covance | 1:1,000 |
| GFP | mouse | Invitrogen | 1:500 |
| MPM2 | mouse | Millipore | 1:200 |
| N-Cadherin | mouse | BD bioscience | 1:100 |
| aPKC (PKC$\zeta$) | rabbit | Santa Cruz | 1:100 |
| GFP | chicken | Aves | 1:500 |
| c-Myc (detecting tdTomato-c-Myc) | goat | Novus | 1:150 |

- **Secondary antibodies**

| Antibody name | Species | Vendor | Dilution |
|---|---|---|---|
| Alexa Fluor 488-, 568-, 647-conjugated antibodies | Donkey anti-mouse, rabbit, rat | Invitrogen | 1:500 |
| Alexa Fluor 594-, 633-conjugated phalloidin | | Invitrogen | 1:100 |
| Anti-GFP and c-Myc | Donkey anti-chicken, goat | Jackson Immuno Research Lab | 1:100 |

- **Retrovirus plasmid construction**

| Retrovirus plasmid | Template plasmid | Source | Other |
|---|---|---|---|
| pCX-mCherry-$\alpha$Tub | pEGFP-$\alpha$Tubulin<br>pRSET-mCherry | Previously generated.<br>Moon et al., 2014 | |
| pCX-MRLC1-GFP | pEGFP-MRLC1 | Tom Egelhoff (Cleveland Clinic, Cleveland, OH, USA) | Addgene plasmid #35680 |
| pCX-SEPT6-GFP | pEGFP-SEPT6 | Matthew Krummel (UCSF, San Francisco, CA, USA) | Described in Gilden et al., 2012 |
| pCX-H2B-tdTomato | pCLNR-H2BG | Geoffrey Wahl (Salk Institute, San Diego, CA, USA) | Addgene plasmid #17735 |
| | pRSET-B-tdTomato | Roger Tsien (UCSD, San Diego, CA, USA) | |
| pCX-DN-RhoA,<br>pCX-CA-RhoA | pEGFP-DN-RhoA (T19N)<br>pEGFP-CA-RhoA (Q63L) | Kozo Kaibuchi (Nagoya University, Nagoya, Japan) | |
