## Supplementary Movie Legend for "LIS1 determines cleavage plane positioning by regulating actomyosin-mediated cell membrane contractility"

**SUPPLEMENTARY MATERIALS**

**Movie S1. Cytokinesis defects in *Lis1* mutant MEFs.**
Time-lapse live cell imaging of mitotic cell division from *Lis1* mutant MEFs ($CreER^{TM}$; $Lis1^{hc/hc}$ +$TM^{12\ h}$) treated with 4-hydroxy tamoxifen (TM). H2B-GFP and mCherry-α-Tubulin labeled fluorescence signals were acquired with a 1 minute interval by Nikon Ti epifluorescence microscope. During cytokinesis of *Lis1*-deficienct MEFs, severe cytokinesis defects were observed such as vigorous cell shape oscillation and spindle rocking.

**Movie S2. Formation of binucleated daughter cells by cytokinesis failure in *Lis1* mutant MEFs.**
Time-lapse live cell imaging of mitotic cell division from *Lis1* mutant MEFs ($CreER^{TM}$; $Lis1^{hc/hc}$ +$TM^{12\ h}$) treated with 4-hydroxy TM. H2B-GFP and mCherry-α-Tubulin labeled fluorescence signals were acquired with a 30 second interval by Nikon Ti epifluorescence microscope. The *Lis1*-deficienct MEFs underwent abnormal cytokinesis and resulted in formation of binucleated daughter cells.

**Movie S3. Myosin II localization during cytokinesis of wild-type (WT) MEFs.**
Time-lapse live cell imaging of mitotic cell division from wild-type (WT) MEFs. Myosin regulatory light chain 1 (MRLC1)-GFP and H2B-tdTomato labeled fluorescence signals were acquired with a 30 second interval by Nikon Ti spinning disk confocal microscope. In normal cytokinesis, MRLC1 was recruited to the equatorial cortex and formed the cleavage furrow.

**Movie S4. Abnormal Myosin II movements during cytokinesis of *Lis1* mutant MEFs.**
Time-lapse live cell imaging of mitotic cell division from *Lis1* mutant MEFs ($CreER^{TM}$; $Lis1^{hc/hc}$ +$TM^{24\ h}$). Myosin regulatory light chain 1 (MRLC1)-GFP and H2B-tdTomato labeled fluorescence signals were acquired with a 30 second interval by Nikon Ti spinning disk confocal microscope. In *Lis1* mutant MEFs, MRLC1 was first recruited to the equatorial cortex. However, we found failure to properly restrict the cleavage furrow at the equatorial cortex.

**Movie S5. Uncoupling between chromosome segregation and cytokinesis in *Lis1* mutant MEFs.**
Time-lapse live cell imaging of mitotic cell division from *Lis1* mutant MEFs ($CreER^{TM}$; $Lis1^{hc/hc}$ +$TM^{24\ h}$). Myosin regulatory light chain 1 (MRLC1)-GFP and H2B-tdTomato labeled fluorescence signals were acquired with a 30 second interval by Nikon Ti spinning disk confocal microscope. We observed frequent uncoupling between chromosome segregation and cytokinesis in *Lis1* mutant MEFs.

**Movie S6. Septin localization during cytokinesis of wild-type (WT) MEFs.**
Time-lapse live cell imaging of mitotic cell division from wild-type (WT) MEFs. Septin 6 (SEPT6)-GFP labeled fluorescence signals were acquired with a 30 second interval by Nikon Ti spinning disk confocal microscope. In normal cytokinesis, Septin-associated contractile ring complex was recruited to the equatorial cortex and formed the cleavage furrow.

**Movie S7. Abnormal Septin localization during cytokinesis of *Lis1* mutant MEFs.**
Time-lapse live cell imaging of mitotic cell division from *Lis1* mutant MEFs ($CreER^{TM}$; $Lis1^{hc/hc}$ +$TM^{24\ h}$). Septin 6 (SEPT6)-GFP labeled fluorescence signals were acquired with a 30 second interval by Nikon Ti spinning disk confocal microscope. We observed SEPT signals at the equatorial cortex initially but then it regressed with vigorous cortical deformation and chromosome oscillation/rocking.
